# The ketone body acetoacetate activates human neutrophils through FFA2R

**DOI:** 10.1101/2022.12.30.522309

**Authors:** Jonas Mårtensson, Lena Björkman, Simon Lind, Moa Bjerhem Viklund, Linjie Zhang, Saray Gutierrez, Claes Dahlgren, Martina Sundqvist, Xin Xie, Huamei Forsman

## Abstract

Neutrophils express many surface receptors that sense environmental changes. One such sensor is FFA2R (free fatty acid receptor 2), a receptor that detects gut microbiota-derived short chain fatty acids. As such, FFA2R has been regarded as a molecular link between metabolism and inflammation. Our recent studies on FFA2R, using its endogenous agonist propionate in combination with allosteric modulators, have identified several novel aspects of FFA2R regulation. A recent study has also identified the ketone body acetoacetate as an endogenous ligand for mouse FFA2R. Whether human FFA2R also recognizes acetoacetate and how this recognition modulates human neutrophil functions has not been earlier investigated. In this study, we found that acetoacetate can induce a decrease of cAMP and translocation of β-arrestin in cells overexpressing FFAR2. In addition, we show that similar to propionate, FFA2R specific allosteric modulators enhance acetoacetate-induced transient rise in cytosolic calcium, production of reactive oxygen species and cell migration in human neutrophils. In summary, we demonstrate that human neutrophils recognize the ketone body acetoacetate through FFA2R. Thus, our data further highlight the key role of FFA2R in inflammation and metabolism.

## Introduction

Neutrophils are the most abundant leukocyte in human blood and perform crucial tasks in the immune defense against invading pathogens. Two main aspects are essential for an effective immune defense by neutrophils: first, the recognition of specific danger- and pathogen-associated molecular patterns (DAMPs and PAMPs) and other markers for environmental changes via specific neutrophil surface receptors, and second, displaying an ample array of weapons to successfully neutralize and eliminate a threat [1]. Many PAMPs and DAMPs are recognized by membrane-bound receptors belonging to the family of G protein-coupled receptors (GPCRs) and the short chain fatty acid receptor FFA2R [2, 3] is one of these.

This receptor (earlier called GPR43) was deorphanized as the receptor for the short chain fatty acids (SCFAs) acetate, propionate and butyrate that are produced by gut bacteria during fiber fermentation [4, 5]. FFA2R is a member of the GPCRs subfamily of free fatty acid receptors (FFARs) sensing fatty acids of different length [6] and it has been highlighted as a potential pharmacological target for treating both inflammatory and metabolic diseases [7, 8]. Our recent studies have revealed several molecular insights into allosteric modulation and activation mechanisms in human blood neutrophils [9–11]. For example, our data show that FFA2R display multiple allosteric binding sites that are structurally separated from each other as well as from the binding site for orthosteric agonists [9]. In addition, we have shown that activation signals transduced by the allosterically modulated FFA2R is biased/functional selective, and that the modulated receptor may be transactivated by signals generated by other GPCRs expressed in neutrophils [9, 10].

In addition to SCFAs, a recent study has identified the ketone body acetoacetate as a ligand for mouse FFA2R [12]. The ketone metabolites, acetoacetate and β-hydroxybutyrate (BHB), are produced in the liver through fatty acid oxidation in response to low glucose levels in blood [13]. When formed, the ketones serve not only as an alternative fuel source during hypoglycemic conditions such as starvation or intense exercise, but also as signaling molecules [14, 15]. A recent study has linked ketone bodies, in particular BHB, to inflammation. This study shows that BHB directly inhibits activation of the NLRP3 inflammasome, resulting in a reduced release of the pro-inflammatory cytokines IL-1β and IL-18 [16]. Recent studies have also shown that increased levels of ketone bodies have beneficial effect for inflammatory conditions including colitis and gout [17, 18]. Nevertheless, despite the increased scientific interest and widespread use of ketogenic diets or carbohydrate-restricted diets [19], very little is known about the impact of the endogenously produced ketone metabolite acetoacetate in modulating inflammation at the molecular level.

In the current study, we explored the ability of human FFA2R to sense acetoacetate and the impact of this recognition in regulating human neutrophil functions. Our data show that human neutrophils selectively sense the ketone body acetoacetate but not BHB and that the recognition is mediated through FFA2R. The activation profile and signals generated by acetoacetate downstream of FFA2R are very similar to those induced by short chain fatty acids. In the presence of positive allosteric FFA2R modulators, acetoacetate acts as a neutrophil chemoattractant and triggers superoxide production. In summary, we demonstrate that human FFA2R senses not only short chain fatty acids but also acetoacetate, and that this ketone body activates human neutrophils and potentially modulate the inflammatory response.

## Material and Methods

### Ethics Statement

This study includes buffy coats/blood obtained from the blood bank at Sahlgrenska University Hospital, Gothenburg, Sweden using blood from healthy human blood donors. According to the Swedish legislation section code 4§ 3p SFS 2003:460, no ethical approval was needed since the buffy coats/blood were provided anonymously.

### Chemicals

The tripeptide fMLF, PMA, propionate, isoluminol, luminol, TNF-α, hexadecyltrimethylammonium bromide (CTAB), bovine serum albumin (BSA), horseradish peroxidase (HRP), *R*-3-hydroxybutyric acid (denoted as BHB) and Lithium-acetoacetate were obtained from Sigma (Sigma Chemical Co., St. Louis, MO, USA). Acetoacetic sodium salt was from MedChemExpress (Monmouth Junction, NJ, USA). Of note, Lithium-acetoacetate was used for all experiments except the cAMP assay where Acetoacetic sodium salt was used. Dextran was from Pharmacosmos (Holbaek, Denmark) and Ficoll-Paque was obtained from GE-Healthcare Bio-Science (Uppsala, Sweden). Fura-2 AM was from Life Technologies (Carlsbad, CA, USA). Superoxide dismutase (SOD) and catalase were obtained from Worthington (Lakewood, NJ, USA). Sytox Green and Lipofectamin 3000 was obtained from Invitrogen (Waltham, MA, USA). The phenylacetamide compound (S)-2-(4-chlorophenyl)-3,3-dimethyl-N-(5-phenylthiazol-2-yl)butanamide, abbreviated as Cmp58, was obtained from Calbiochem (San Diego, CA, USA). AZ1729 was a kind gift from AstraZeneca (Gothenburg site, Sweden). The FFA2R antagonist CATPB ((S)-3-(2-(3-chlorophenyl)acetamido)-4-(4-(trifluoromethyl)-phenyl) butanoic acid) was from Tocris (Britsol, UK).

Subsequent dilutions of all reagents were made in Krebs-Ringer Glucose phosphate buffer (KRG; 120 mM NaCl, 4.9 mM KCl, 1.7 mM KH_2_PO_4_, 8.3 mM NaH_2_PO_4_, 1.2 mM MgSO_4_, 10 mM glucose, and 1 mM CaCl_2_ in dH_2_O, pH 7.3).

### FFA2R-mediated cyclic adenosine monophosphate (cAMP) assay

HEK293 cells were transfected with plasmids encoding human FFA2R 24 hours before the assay. Cells were harvested and re-suspended in Dulbecco’s modified eagle medium (DMEM) containing 500 μM IBMX (an inhibitor of cyclic nucleotide phosphodiesterase) at a density of 4 × 10^5^ cells/ml. Cells were then plated onto 384-well assay plate at 2000 cells/5 μl/well. Another 5 μl buffer containing compounds at various concentrations were added to the cells. After incubation at room temperature for 30 minutes, 5 μl DMEM containing 1 μM forskolin (to increase the level of cAMP so the subsequent lowering of the cAMP level due to Gαi activation can be observed) was added, and the incubation was continued for another 30 minutes. Intracellular cAMP measurement was carried with a LANCE Ultra cAMP kit (PerkinElmer, Cat No: TRF0264, Waltham, USA) using an EnVision multiplate reader (PerkinElmer, Waltham, USA) according to the manufacturer’s instructions. This assay is a homogenous time-resolved fluorescence resonance energy transfer (TR-FRET) immunoassay, based on the competition between the europium (Eu) chelate-labeled cAMP tracer and cellular cAMP for binding sites on cAMP-specific monoclonal antibodies. A decrease in intracellular cAMP results in higher FRET signal.

### FFA2R-mediated β-arrestin2 recruitment

The recruitment of β-arrestin2 to FFA2R was measured using the Promega NanoBiT Protein-Protein Interaction System [20]. In brief, HEK293 cells (2 × 10^6^) were transfected with plasmids encoding SmBiT-β-arrestin2 (0.5 μg) and FFA2R-LgBiT (0.5 μg) by Lipofectamin 3000. The cells were seeded at 3.6 ×10^4^ cells/well on 96-well plates and incubated at 37°C, 5% CO_2_. The cell media were replaced 48 hours later with 40 μl Opti-MEM media. Thereafter 10 μl Nano-Glo Live Cell reagent was added according to the manufacturer’s protocol (Promega, Cat No: N2011, Madison, USA), and the cells were incubated at 37°C, 5% CO_2_, for five minutes before another 25 μl Opti-MEM media with various concentrations of compounds were added to the cells. Bioluminescence was measured immediately after stimulation for twenty minutes with an EnVision multiplate reader (PerkinElmer, Waltham, USA).

### Isolation of neutrophils from human blood

Neutrophils were isolated from buffy coats or from whole blood from healthy blood donors by dextran sedimentation and Ficoll-Paque gradient centrifugation as described by Bøyum [21]. Remaining erythrocytes were removed by hypotonic lysis and the neutrophils were washed twice and resuspended in KRG and stored on ice until use.

### Calcium mobilization

Neutrophils at a density of 1–3×10^6^ cells/ml were washed with Ca^2+^-free KRG and centrifuged at 220x*g*. The cell pellets were re-suspended at a density of 2×10^7^ cells/ml in KRG containing 0.1% BSA, and loaded with 2 μM Fura 2-AM for 30 minutes at room temperature. The cells were then washed once with KRG and resuspended in the same buffer at a density of 2×10^7^/ml. Calcium measurements were carried out in a Perkin Elmer fluorescence spectrophotometer (LC50), with excitation wavelengths of 340 nm and 380 nm, an emission wavelength of 509 nm, and slit widths of 5 nm and 10 nm, respectively. The transient rise in intracellular calcium is presented as the ratio of fluorescence intensities (340 nm: 380 nm) detected.

### Measuring extracellular and intracellular NADPH-oxidase activity

Isoluminol/luminol-enhanced chemiluminescence (CL) technique was used to measure superoxide production, the precursor of reactive oxygen species (ROS), by the NADPH-oxidase activity as described [22]. The CL measurements were performed in a six-channel Biolumat LB 9505 (Berthold Co., Wildbad, Germany), using 4-ml disposable polypropylene tubes and a 900 μl reaction mixture containing 10^5^ cells. Cell impermeable isoluminol (2 x 10^-5^ M) and HRP (4 U/mL) was used for determination of extracellular ROS. For intracellular ROS measurement, isoluminol and HRP were exchanged for luminol (2 x 10^-5^ M) and superoxide dismutase (SOD, 50 U/mL) and catalase (2000 U/mL) were added to the reaction mixture to quench extracellular ROS. The tubes were equilibrated for 5 minutes at 37°C, before addition of stimuli (100 μl) and light emission was recorded continuously over time. In experiments where the effects of receptor specific positive allosteric modulators and the receptor specific antagonist CATPB was determined, the compounds were added to the reaction mixture just prior the five minutes equilibration phase. Control cells were incubated under the same conditions but received no modulator/inhibitor. For determination of both extra- and intracellular ROS-production neutrophils were pretreated with the priming agent TNFα (10 ng/ml) and incubated at 37°C for 20 minutes, thereafter stored on ice until use. The light emission/superoxide anion production is expressed as Mega counts per minute (Mcpm).

### NETs formation and DNA Release Measurements with Sytox Green

Neutrophils (5× 10^4^ cells/well) suspended in RPMI (without phenol red) and the Sytox Green DNA stain (1.25 μM) were added to black 96-well plates and incubated at 37°C, 5% CO_2_. Different stimuli were added, and Sytox Green fluorescence was measured at indicated time points at 485/535 nm in a CLARIOstar plate reader (BMG Labtech) as described [23].

### Chemotaxis assay

Freshly prepared neutrophils from whole blood (6×10^5^ suspended in 30 μl buffer) were added on top of the membrane of a cell migration system (ChemoTx chemotaxis system, Neuro Probe, MD, USA) whereas stimuli were added below the membrane to create a gradient as described [24]. Allosteric modulators (Cmp58/AZ1729) where added both on top and below the membrane when used. The membrane and liquid loaded at the top of the membrane were carefully removed after incubation at 37°C, 5% CO_2_, for 90 minutes and 5 μl lysis buffer (PBS containing BSA (2%) and CTAB (2%)) were added to the remaining liquid. 20 μl lysis product was collected after 1 h and transferred to a 96-well plate containing 80 μl peroxidase reagent (2 OPD tablets dissolved in 10 mL TMB-buffer with 4 μl (30%) H_2_O_2_). The MPO content from the samples were determined after 30 min using a plate reader (CLARIOstar (BMG Labtech), absorbance at 450 nm).

### Data analysis

Data analysis was performed using GraphPad Prism 9.3.1 (Graphpad Software, San Diego, CA, USA). Curve fitting was performed by non-linear regression using the sigmoidal dose-response equation (variable-slope). Statistical analysis was performed using a ratio paired *t*-test or a repeated measures one-way ANOVA followed by Dunnett’s multiple comparison test. All statistical analysis was performed on unprocessed raw data and statistically significant differences are indicated by \**p* < 0.05, \*\**p* < 0.01.

## Results

### The ketone acetoacetate is an endogenous agonist for the short chain fatty acid receptor FFA2R

The ketone body acetoacetate was recently identified through ligand screening to be an agonist for mouse FFA2R [12]. Provided that also primary human cells expressing FFA2R can mediate biological functions by ketone metabolites, this adds to the understanding of the signaling properties of ketone bodies and opens for a new research area [25]. To determine the ligand recognition profile of the human FFA2R, we used HEK293 cells overexpressing this receptor. Similar to propionate, acetoacetate dose-dependently activated the receptor determined as an inhibition of forskolin-induced accumulation of intracellular cAMP (Fig 1A, B). In comparison to propionate, acetoacetate was however less potent with an EC_50_ of ~ 1 mM (Fig 1A, B). In agreement with the results shown for mouse FFA2R [12], no activation was induced by the other predominant ketone body β-hydroxybutyrate (BHB; Fig 1C). In a β-arrestin recruitment assay, acetoacetate induced a rapid β-arrestin recruitment mediated by FFA2R, and the recruitment was comparable to that induced by propionate (Fig 1D, E). Even at concentrations up to 30 mM, BHB did not induce any detectable β-arrestin recruitment (Fig 1E). Taken together, these data show that the human FFA2R is a receptor not only for short chain fatty acids but also for the ketone body acetoacetate.

**Fig 1.**
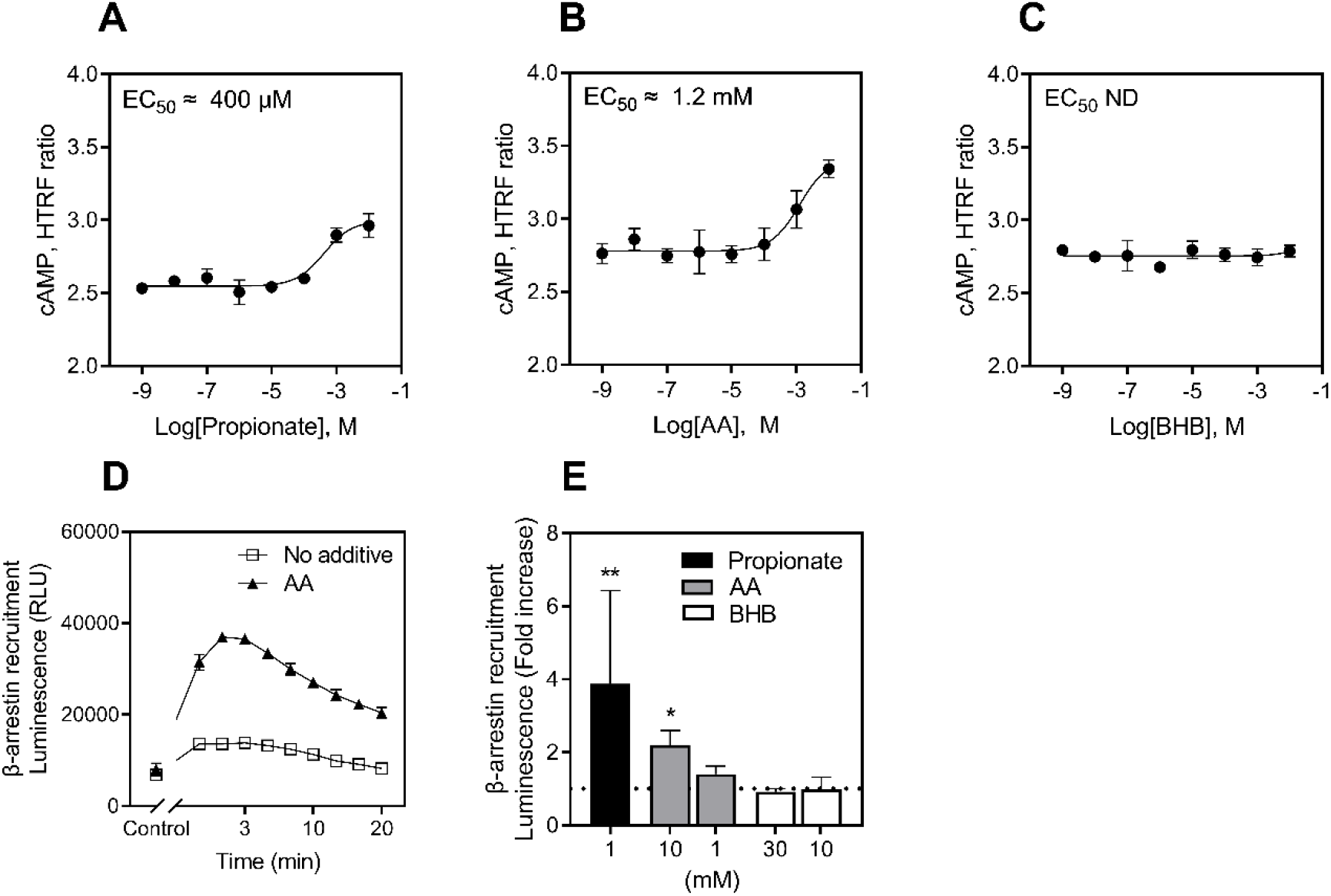
Acetoacetate but not BHB activates FFA2R in HEK293 cells overexpressing the receptor. HEK293 cells overexpressing FFA2R were activated with different concentrations of **A)** propionate, **B)** acetoacetate (AA), and **C)** β-hydroxybutyrate (BHB) and cAMP was measured. Data obtained was fitted to dose-response curves (mean +/− SD, n = 3 independent experiments) and EC_50_ values were calculated. EC_50_ for propionate was determined to 410 μM (95% CI: 199 - 827 μM) and for AA to 1.24 mM (95% CI: 0.45 mM - 3.60 mM). The EC_50_ value of BHB was not possible to determine since no change in cAMP levels were detected (ND = not determined). **D)** β-arrestin translocation was measured continuously over time upon stimulation with acetoacetate 10 mM (closed triangles) and compared to background activity when no stimulus was added (open squares). The control shows the level of luminescence before addition of both stimuli and Nano-Glo reagent. **E)** Bar graph showing the fold increase of peak values, compared to time matched background activity, upon stimulation with propionate (black bar), AA (grey bars) and BHB (white bars). The dotted line indicates the level of background activity. Data are presented as a mean + SD (n= 3 individual experiments) and statistical analysis was performed using ratio paired *t*-test (* = p<0.05, ** = p<0.01).

### Acetoacetate triggers an activation of FFA2R in primary blood neutrophils

A transient change in the concentration of intracellular free calcium ions ([Ca^2+^]_i_,), is an early signaling event down-stream of many activated neutrophil GPCRs including FFA2R [10, 11]. Accordingly, propionate induced a transient change in the [Ca^2+^]_i_ in neutrophils (Fig 2A). Similar to propionate, acetoacetate also induced a dose-dependent transient rise in [Ca^2+^]_i_ in neutrophils (Fig 2B). To determine if the response was mediated through FFA2R the receptor specific antagonist CATPB was introduced. The acetoacetate-induced rise in [Ca^2+^]_i_ could be completely abolished in the presence of CATPB (Fig 2C). In line with our earlier data on the effects of positive allosteric FFA2R modulation [26], the allosteric FFA2R modulator Cmp58, that lacks effect on its own, enhanced the propionate response (Fig 2D). Similarly, Cmp58 enhanced also the acetoacetate response at a concentration that alone was insufficient to induce a rise in [Ca^2+^]_i_ (Fig 2E). These data show that human FFA2R recognizes acetoacetate and a positive allosteric FFA2R modulator enhances the outcome of this recognition. In contrast to acetoacetate, no rise in [Ca^2+^]_i_ was induced by BHB with or without Cmp58 (Fig 2F). Taken together, we show that the ketone body acetoacetate is recognized by FFA2R in human neutrophils, this recognition initiates the signaling cascade leading to a rise of [Ca^2+^]_i_.

**Fig 2.**
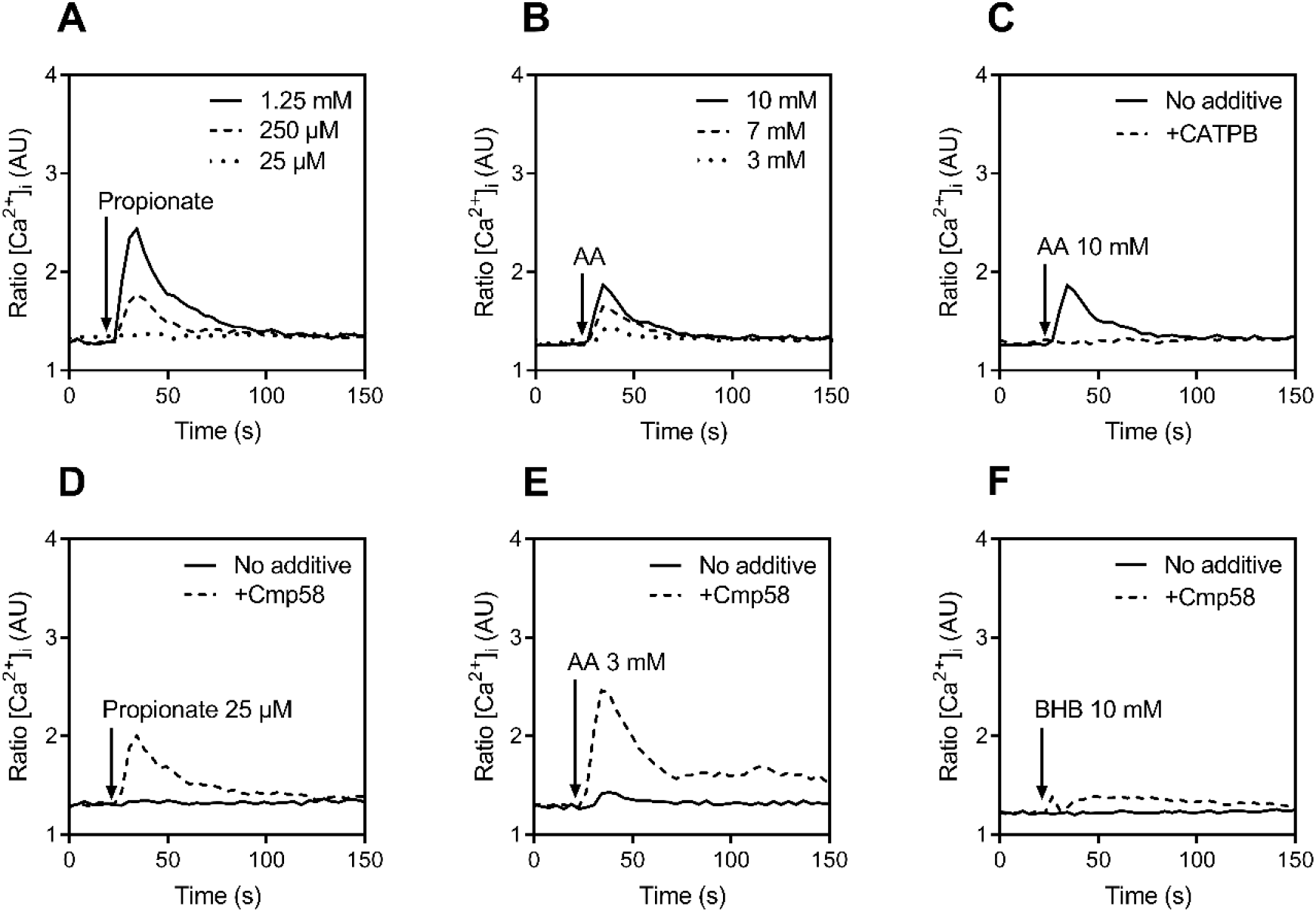
Acetoacetate triggers a transient rise in intracellular calcium ([Ca^2+^]_i_,) in human neutrophils. **A-B)** Human neutrophils were loaded with Fura-2 AM and stimulated with different concentrations of **A)** propionate or **B)** acetoacetate (AA). **C)** Neutrophils were incubated with (dashed line) or without (solid line) the FFA2R antagonist CATPB (100 nM) and then stimulated with AA (10 mM). **D-F)** Neutrophils were incubated with (dashed line) and without (solid line) Cmp58 (1 μM) for 10 minutes and then stimulated with **D)** propionate (25 μM) or **E)** AA (3 mM) or **F)** β-hydroxybutyrate (BHB; 10 mM). The rise in intracellular Ca^2+^ [Ca^2+^]_i_ is presented as the ratio between Fura-2 fluorescence at 340 and 380 nm over time and one representative experiment out of three independent experiments is shown.

### Acetoacetate triggers an FFA2R-dependent secretion of reactive oxygen species (ROS)

Our earlier studies on the effects of propionate and synthetic FFA2R agonists have demonstrated several novel insights into allosteric FFA2R modulation in human neutrophils [9, 10, 26, 27]. Accordingly, in the presence of the allosteric FFA2R modulator Cmp58, we observed that not only propionate but also acetoacetate induced secretion of ROS generated by the neutrophil NADPH-oxidase (Fig 3A-C). None of the agonists induced any ROS production without Cmp58 (Fig 3A-C). In the presence of Cmp58, acetoacetate dose-dependently induced ROS secretion with an EC_50_ of 700 μM (Fig 3D). The response induced by acetoacetate was fully inhibited by the FFA2R specific antagonist CATPB (Fig 3E, F), further confirming that FFA2R is the receptor in human neutrophils not only for short chain fatty acids but also for the ketone body acetoacetate.

**Fig 3.**
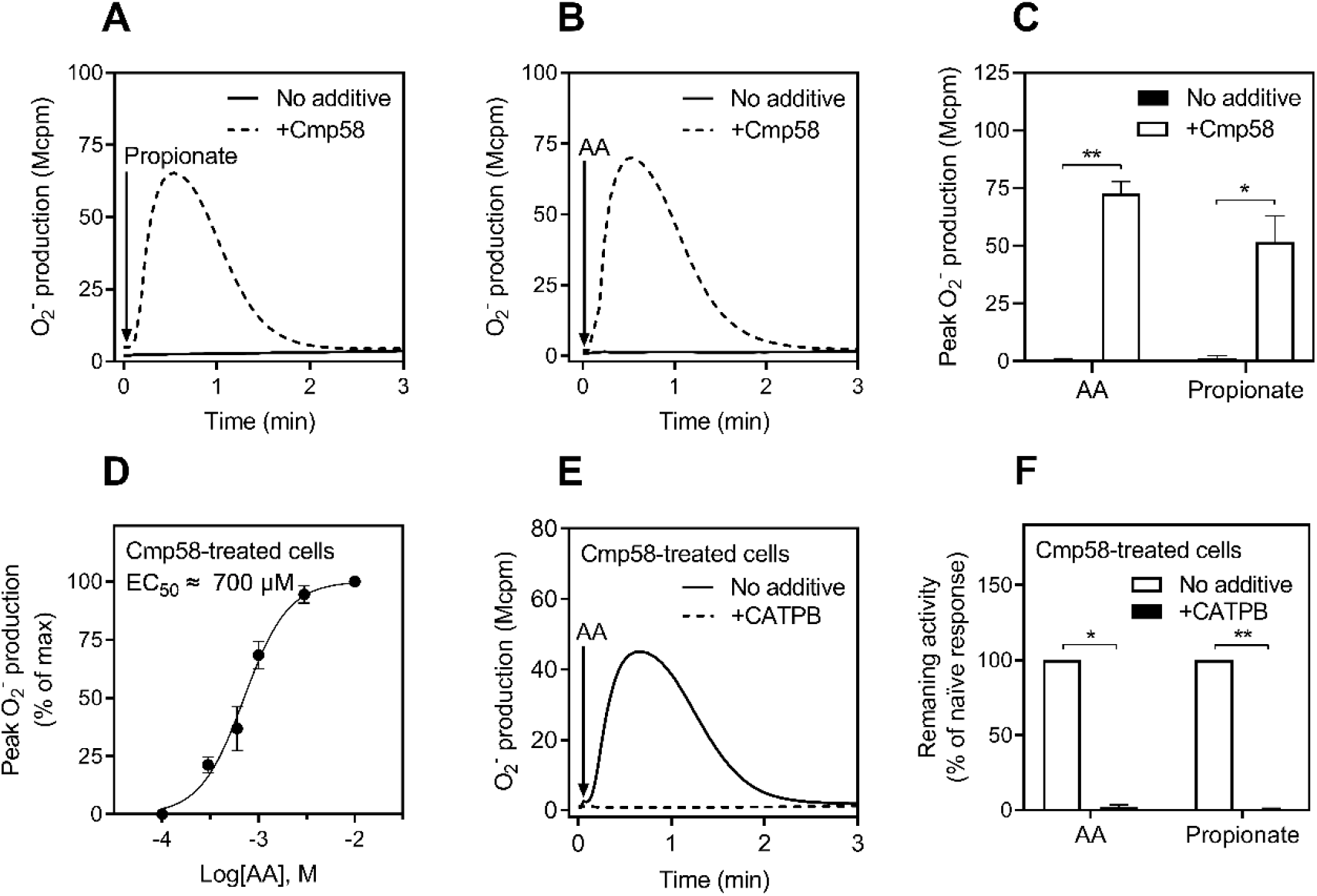
Acetoacetate triggers production of oxygen radicals in neutrophils allosterically modulated by Cmp58. **A-B)** Neutrophils pre-incubated for five minutes in the absence (solid line) or presence (dashed line) of the FFA2R allosteric modulator Cmp58 (1 μM) was stimulated with **A)** propionate (25 μM) or **B)** acetoacetate (AA, 10 mM) and superoxide release was measured over time. **C)** Summary of the peak superoxide production (mean + SD, n = 3 independent experiments) in response to stimulation with AA (10 mM) and propionate (25 μM) in cells treated with (white bars) or without (black bars) Cmp58 (1 μM). **D)** Neutrophils treated with Cmp58 (1 μM) was stimulated with different concentration of AA and superoxide production was recorded. The peak values were determined, fitted to a dose response curve (mean + SD, n = 3 independent experiments) and the EC_50_-value was calculated. **E)** Neutrophils were incubated for five minutes with Cmp58 (1 μM) together with (dashed line) or without (solid line) the FFA2R antagonist CATPB (100 nM) prior to stimulation with AA (1 mM). **F)** Bar graph (mean + SD, n = 3 independent experiments) showing the peak superoxide production in cells incubated with Cmp58 (1 uM) together with (black bars) or without (white bars) CATPB (100 nM) and then stimulated with AA (1 mM) or propionate (25 μM). Statistical analysis in **C** and **F** was performed using a ratio paired *t*-test (* = p<0.05; \*\**p*<0.01).

In summary, we show that no activation of the neutrophil NADPH-oxidase is induced by acetoacetate alone, but in the presence of an allosteric FFA2R modulator, acetoacetate is a strong ROS inducer of this neutrophil enzyme system. This is an activation pattern that acetoacetate share with the short chain fatty acids.

### Allosteric FFA2R modulation and homologous desensitization

To gain more insights into acetoacetate-induced FFA2R activation in human neutrophils, we used different FFA2R modulation models that applies to propionate. One such model is the reciprocal allosteric modulation, *i.e*., similar cellular response is induced when the order of addition of the allosteric modulator Cmp58 and the endogenous agonist propionate was reversed. Similar to this, the order by which Cmp58 and acetoacetate was added could also be reversed (Fig 4A, B). The presence of acetoacetate was clearly required in order for Cmp58 to trigger an activation of the neutrophil superoxide generating system as no ROS was induced by Cmp58 alone (Fig 4B). To further elucidate the requirement of an allosteric FFA2R modulator in acetoacetate-induced neutrophil ROS production, we replaced Cmp58 with AZ1729. AZ1729 is another positive allosteric FFA2R modulator earlier shown to bind an allosteric FFA2R site, different from that of Cmp58, and to modulate neutrophil functions similarly to Cmp58 [9]. The neutrophil response induced by acetoacetate was positively modulated also by AZ1729, as shown by a robust production of ROS (Fig 4C, D). These results further strengthen the conclusion that acetoacetate is an FFA2R agonist and a strong inducer of ROS for human neutrophils provided that FFA2R is allosterically modulated.

**Fig 4.**
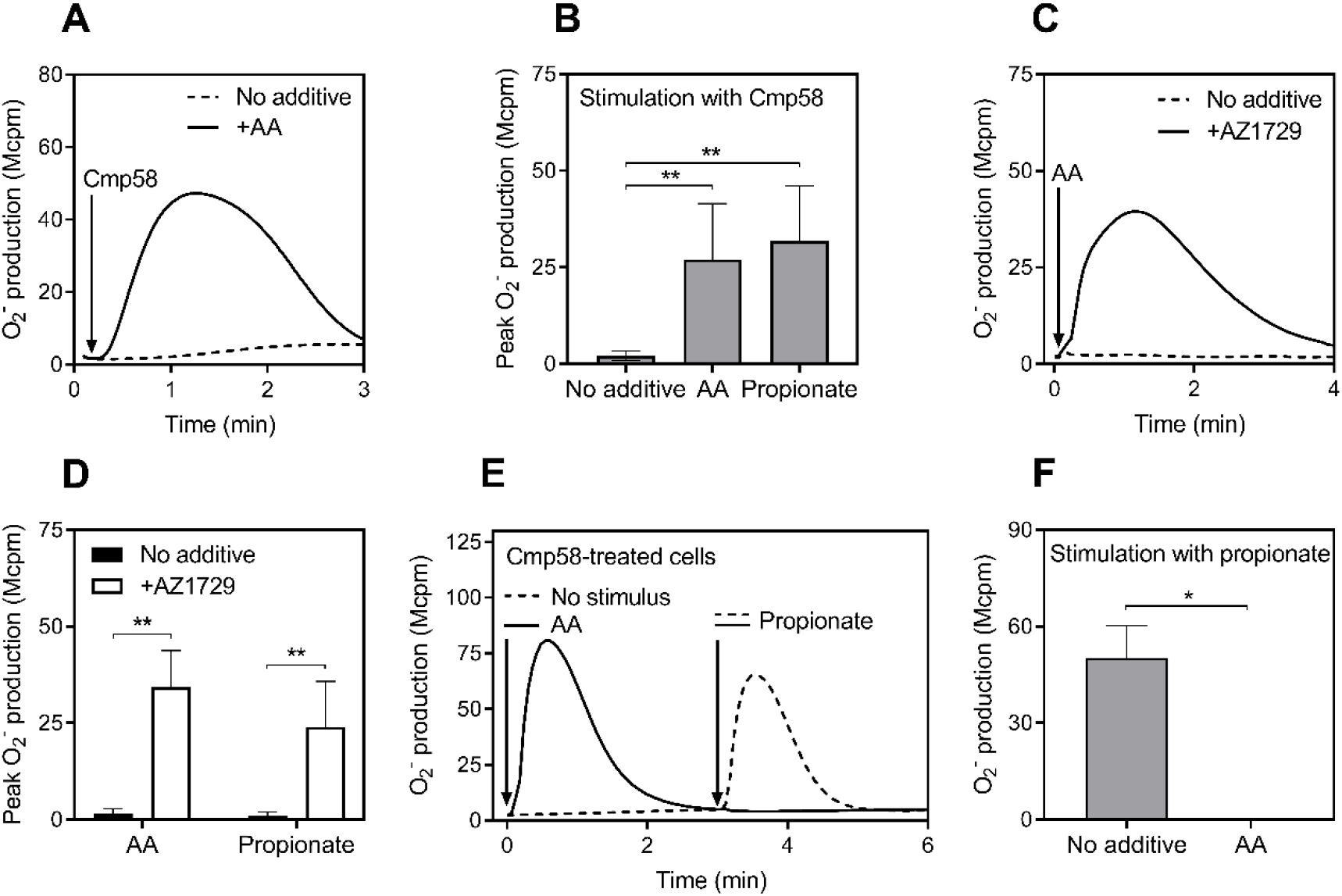
Characterization of the interaction between acetoacetate and FFA2R. **A)** Superoxide production by neutrophils pre-incubated for five minutes in the absence (dashed line) or presence (solid line) of acetoacetate (AA; 1 mM) before stimulation with Cmp58 (1 μM) and measurement of superoxide release over time. **B)** Summary of the peak superoxide production (mean + SD, n = 4 independent experiments) in response to stimulation with Cmp58 (1 μM) in cells pre-treated with AA (1 mM) or propionate (25 μM) compared to the naïve Cmp58 response. **C)** Superoxide production by neutrophils pre-incubated for five minutes in the absence (dashed line) or presence (solid line) of AZ1729 (1 μM) before stimulation with AA (10 mM) and measurement of superoxide release over time. **D)** The bar graph (mean + SD, n = 3 independent experiments) show the peak value of superoxide production by neutrophils pre-treated for five minutes with (white bars) or without (black bars) AZ1729 (1 μM) prior stimulation with AA (10 mM) or propionate (25 μM). **E)** Neutrophils incubated with Cmp58 (1 μM) for five minutes were stimulated with either AA (10 mM, solid line) or buffer (dashed line) at the time point indicated by the left arrow. When the response had returned to basal levels the neutrophils were re-stimulated with propionate (25 μM; time of addition shown by right arrow). **F)** Summary of the peak superoxide production (mean + SD, n = 3 independent experiments) induced by propionate in cells first stimulated with either AA (10 mM) or buffer (no additive). Statistical analysis in **B** was performed using a repeated measures one-way ANOVA followed by Dunnett’s multiple comparison test, and for **D** and **F** a ratio paired *t*-test was performed (* = p<0.05; ** = p<0.01).

Next, we examined the ability of FFA2R to undergo homologous desensitization upon stimulation, *i.e*., receptors become non-responsive to a second stimulation with either the same agonist or another agonist that binds to the same receptor [3]. Accordingly, we found that in the presence of Cmp58, the active signaling state of FFA2Rs when challenged with acetoacetate was rapidly terminated in less than three minutes. When the response to acetoacetate had declined, these neutrophils were non-responsive (desensitized) to a second stimulation with propionate (Fig 4E, F). These data further confirm that acetoacetate and propionate bind to the same receptor and can homologously desensitize FFA2R in human neutrophils.

Taken together, the data presented show that acetoacetate activates neutrophils through FFA2R and regulate receptor activity by similar mechanisms as propionate.

### Acetoacetate triggers an FFA2R-dependent intracellular ROS production but does not induce formation of neutrophil extracellular traps (NETs)

The neutrophil NADPH-oxidase can be assembled and activated not only at the plasma membrane, resulting in a secretion of ROS, but also in granule membranes giving rise to intracellular production of ROS [22, 28]. We examined the ability of acetoacetate to trigger intracellular ROS production. Unlike PMA (a PKC activating phorbol ester used as control compound), acetoacetate alone was unable to induce intracellular ROS production (Fig 5A), however, when combined with Cmp58 also acetoacetate triggered an intracellular production of ROS (Fig 5A, B). This further supports the notion that the neutrophil NADPH oxidase can be assembled intracellularly, resulting in ROS generation within intracellular organelles.

**Fig 5.**
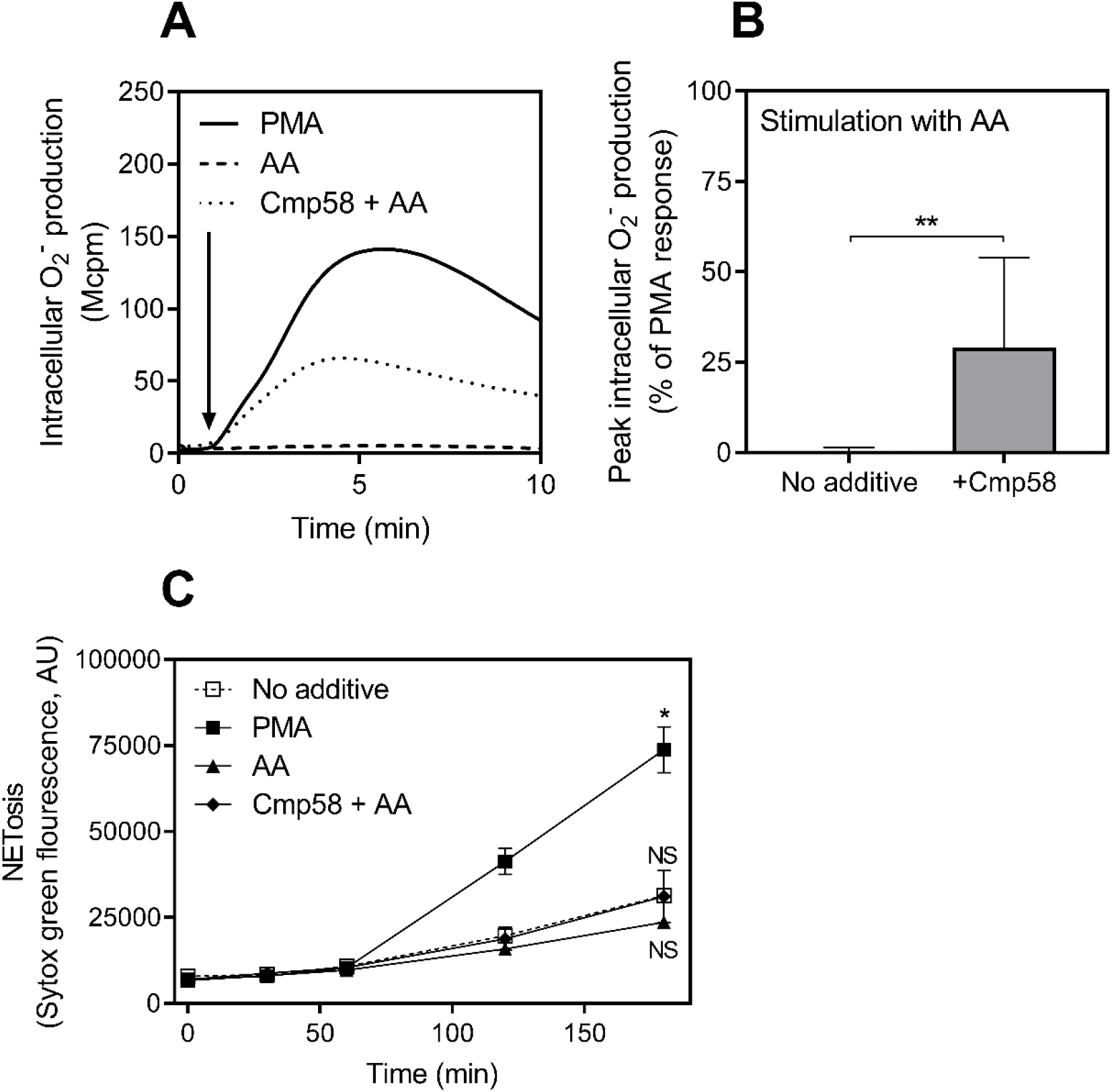
Neutrophils allosterically modulated with Cmp58 produce intracellular radicals but does not trigger NETosis. **A)** Intracellular production of oxygen radicals in response to acetoacetate (AA; 1 mM) was measured over time in neutrophils pre-treated with (dotted line) and without (dashed line) Cmp58 (1 μM) for five minutes. Stimulation with phorbol myristate acetate (PMA; 50 nM) was performed in parallel as a reference (solid line). **B)** Summary of the peak intracellular radical production (mean + SD, n = 5 independent experiments) induced by AA (1 mM) in neutrophils pre-incubated with and without Cmp58 (1 μM) for five minutes. **C)** Neutrophil NETosis (mean + SD; n = 3 independent experiments) was determined by measuring levels of nucleic acid exposed extracellularly (using Sytox green staining) over time in response to stimulation with PMA (50 nM, line with filled squares), AA (10 mM; line with triangles) and AA (10 mM) in presence of Cmp58 (1 μM, line with diamonds). The background level of fluorescence is included as a reference (no additive, line with open squares). Statistical analysis in **B** was performed by a ratio paired *t-test* and for **C** by using a repeated measures one-way ANOVA followed by Dunnett’s multiple comparison test against no additive for values at timepoint 180 minutes (* = p<0.05; ** = p<0.01).

The precise biological role of the intracellular generated ROS in the absence of phagocytosis is not clear, but it has been suggested that ROS mediate signals with the capacity to limit autoimmune and inflammatory reactions [28–30]. In addition, an assembly and production of intracellular ROS has been shown to be essential for formation of neutrophil extracellular traps (NETs) induced by some, but not all stimuli [23, 31]. It has been suggested that SCFAs at intestinal concentrations trigger NET formation [32], and with this as the backdrop we examined whether the ROS produced intracellularly by FFA2R activation contribute to the formation of NETs. PMA stimulation was used as positive control and NET formation was determined by staining of DNA released from the neutrophils (Fig 5C). We found that, despite the ability of acetoacetate to induce intracellular ROS production, no NETs were formed within three hours of monitoring (Fig 5C). Taken together, our data show that acetoacetate triggers FFA2R signaling leading to intracellular ROS production without any formation of NETs.

### Acetoacetate attracts neutrophils in the presence of allosteric FFA2R modulators

Many agonists recognized by neutrophil GPCRs trigger not only an activation of the ROS producing NADPH-oxidase, but they also function as chemoattractants [1, 33]. To determine the ability of acetoacetate to induce neutrophil recruitment, we used a cell migration assay in which neutrophils were allowed to migrate through a filter that separates neutrophils from the recruiting agonist. Compared to the well-known chemoattractant fMLF (an FPR1-agonist), no significant migration was induced by either propionate or acetoacetate alone at all concentrations tested (Fig 6A). However, in the presence of either of the positive FFA2R allosteric modulators Cmp58 or AZ1729, not only propionate but also acetoacetate was chemotactic and induced neutrophil migration to levels similar to that induced by fMLF (Fig 6B-E). In summary, we show that acetoacetate not only induces ROS production but also attracts neutrophils chemotactically, provided that FFAR2 has been allosterically modulated.

**Fig 6.**
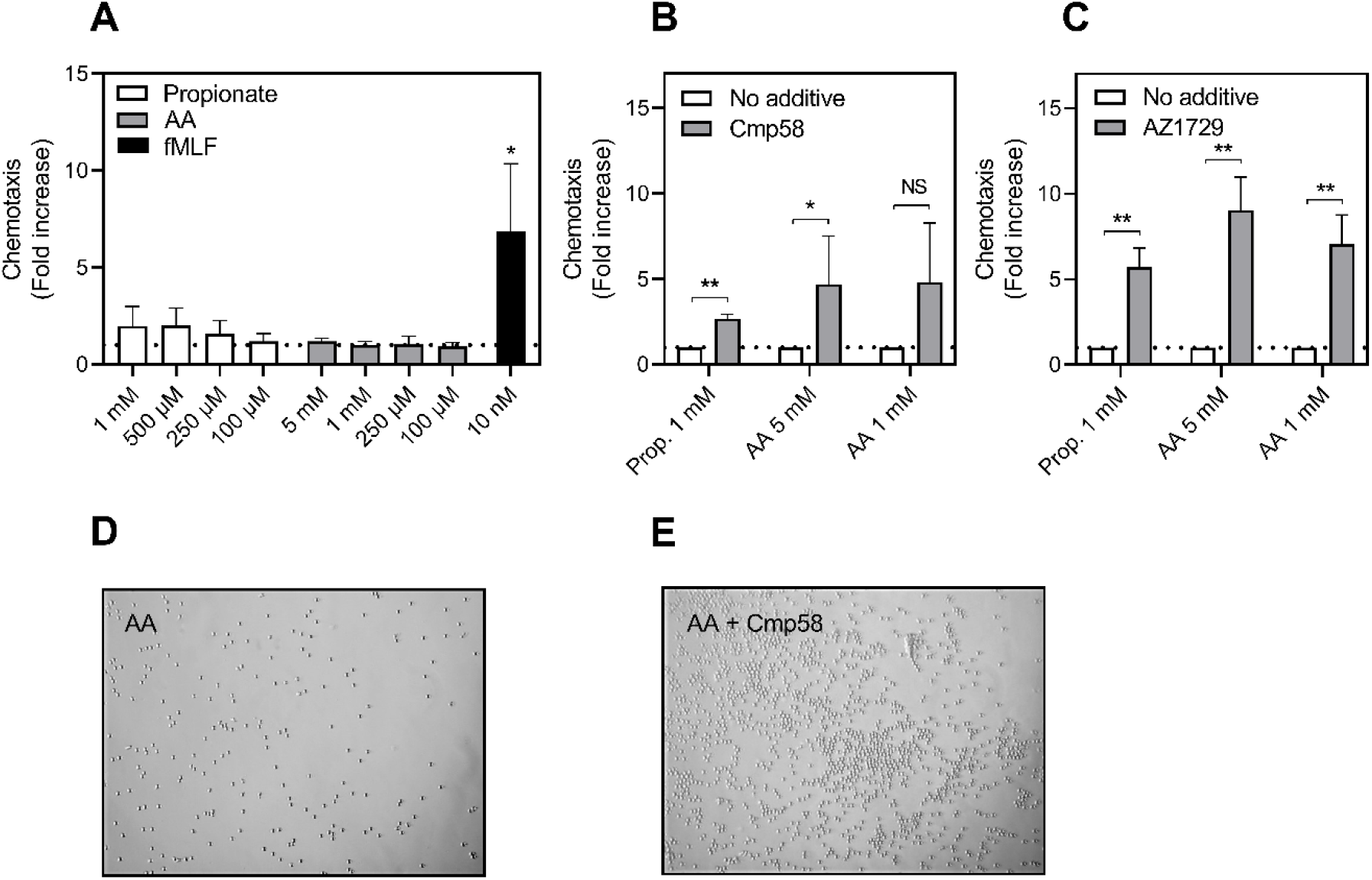
Acetoacetate induces chemotaxis in neutrophils treated with FFA2R-positive allosteric modulators. **A)** Neutrophil chemotaxis (n = 3 independent experiments) induced by different concentrations of propionate (white bars) or acetoacetate (AA; grey bars), fMLF included as a positive control (black bars, 10 nM). Data are presented as fold increase of spontaneous migration towards buffer (level of spontaneous migration is indicated with the dotted line). **B-C)** Neutrophil chemotaxis (n = 3 independent experiments) induced by propionate (1 mM) and AA (5 mM and 1 mM) in the absence (white bars) or presence (grey bars) of either of the allosteric modulators **B)** Cmp58 (1 μM) or **C)** AZ1729 (1 μM). Data are presented as fold increase of migration towards the stimuli in the absence of allosteric modulator (level indicated by the dotted line). **D-E)** Representative microscopic images of the neutrophils after allowing for migration for 90 min towards **D)** AA (5 mM) or **E)** AA (5 mM) in the presence of Cmp58 (1 μM). Statistical analysis in **A** was performed using a repeated measures one-way ANOVA followed by Dunnett’s multiple comparison test against buffer. A ratio paired *t*-test was used for analysis of **B** and **C** (* = p<0.05; ** = p<0.01; NS = no statistically significant difference).

## Discussion

The aim of our study was to investigate whether the ketone body acetoacetate is recognized by human FFA2R as an endogenous ligand and by that has the capacity to modulate the inflammatory process through neutrophil activation. Our data show that human FFA2R recognizes not only short chain fatty acids but also the ketone body acetoacetate. This finding supports the recent attention that FFA2R plays an important role in regulating metabolism and inflammation [6, 7, 34]. In addition, our study provides molecular evidence that the ketone body acetoacetate, shown to be a novel endogenous FFA2R agonist that activates human neutrophils, should be included as a modulator of inflammation. This new knowledge could possibly increase our understanding of how diet and metabolic status alter inflammatory reactivity.

Our data obtained with cells that overexpress human FFA2R show that acetoacetate but not BHB, is an endogenous activating ligand of FFA2R. Our results are in agreement with previous published data on mouse FFA2R and in animal models showing that FFA2R selectively recognizes acetoacetate but not BHB [12]. Ketone bodies (predominantly acetoacetate and BHB) are produced in the liver under hypoglycemic conditions such as prolonged fasting and intense exercise [13]. Acetoacetate and BHB are used as alternative energy sources by skeletal muscle, heart, and brain in conditions of glucose deprivation [15]. However, beyond their use as fuels, new immune functions for ketone bodies have recently been described. For example, studies have shown that BHB has beneficial effects in animal disease models of gout and colitis [17, 18]. BHB has also been shown to regulate inflammation and inhibit inflammasome activation and the effect has been suggested to be mediated through receptors such as the nicotinic acid receptor GPR109A [35]. We now add to the evidence that also acetoacetate can directly modulate inflammation through the activation of human neutrophils and that the responsible receptor is FFA2R which is shared by the gut microbiota-derived short chain fatty acid metabolites. The fact that FFA2R is able to recognize both acetoacetate and SCFAs, suggests that these two metabolites cooperatively regulate FFA2R function. Indeed, it has been shown that the peripheral blood concentrations of two metabolites can switch in mice during fasting; an increase of ketone body concentrations is accompanied with a decrease of SCFAs in the circulation, and that diet shapes the gut microbiota [12, 36]. In addition, the beneficial effect of a ketogenic diet has been reported on conditions ranging from inflammatory conditions, epilepsy, cardiac diseases to cancer [19, 37].

By using two earlier characterized allosteric FFA2R modulators, recognized by two different allosteric sites on FFA2R, we could show that similar to propionate, also acetoacetate-induced neutrophil responses were amplified. In the presence of the allosteric modulators AZ1729 or Cmp58, acetoacetate induced neutrophil migration and ROS-production. In the absence of allosteric modulators, acetoacetate is only a very weak agonist similar to the short chain fatty acids. However, the recognition of acetoacetate by human FFA2R in the absence of allosteric modulators is also evident as illustrated by the fact that acetoacetate alone is able to trigger a decrease in cAMP levels, a transient rise in intracellular Ca^2+^ as well as recruitment of β-arrestin. Although we and others have clearly demonstrated that FFA2R activity is tightly regulated by the allosteric modulators [9, 38], an allosteric modulator of endogenous origin has not yet been described. Accordingly, the role of allosteric modulation of FFA2R *in vivo* can at present only be speculated upon. If allosteric modulation of FFA2R is an *in vivo* phenomenon, this could possibly be explored for therapeutic applications. It is also possible that neutrophil activation through FFA2R is tightly controlled locally at the site of inflammation, *i.e*., a proper activation requires the presence of all three partners (neutrophils, allosteric modulators and endogenous agonists). The latter might be a control mechanism to allow restricted and temporal FFA2R-mediated neutrophil activation in order to avoid unnecessary damage to surrounding tissues. Taken together, our data suggest that regulation of FFA2R activity is tightly controlled, and the neutrophil activation profile is very similar for acetoacetate and SCFAs.

In subsequent experiments, we show that acetoacetate, in the presence of allosteric FFA2R modulators, is a neutrophil chemoattractant. Although the mechanism for recruitment remains to be determined, our earlier data on *in vivo*-transmigrated human neutrophils, suggest that FFA2R has a role in guiding neutrophils to the site of inflammation [39]. Regarding the ROS production induced by acetoacetate through FFA2R, there are many implications in regulation of inflammation. One implication is that intracellular ROS formed in neutrophils through granule-granule fusion has been regarded as a determinant for the formation of neutrophil extracellular traps (NETs) induced by certain stimuli including PMA [23]. Our data show that despite the intracellular production of ROS induced by acetoacetate when combined with the allosteric modulator Cmp58, the amount of ROS (less than half of the intracellular ROS induced by PMA) or production of additional intracellular signals generated downstream of FFA2R are not sufficient to trigger formation of NETs. The signaling cascade induced by FFA2R follows the classic G protein/β-arrestin pathways that is not used by PMA that directly activate PKC [3]. Other implications of ROS in inflammation include their role in cell signaling and suppression of inflammation (for more details, see review [28]). Our *in vitro* data indicates that acetoacetate is of importance for modulation of inflammation through neutrophil FFA2R. Whether this translates to an *in vivo* setting and if the acetoacetate-FFA2R interaction partially explain the beneficial effects observed with a ketogenic diet on range of medical conditions can just be speculated upon. As of today, the precise physiological/pathophysiological role of FFA2R is not fully understood.

In summary, we show that acetoacetate is an endogenous agonist for human FFA2R and provide novel insights into the molecular processes behind acetoacetate modulation of neutrophil function. Our finding adds yet another molecular link between metabolism and inflammation. The emerging evidence of the beneficial effect of a ketogenic state on a wide range of medical conditions calls for more studies in the future to investigate the neutrophil-FFA2R-ketone body axis.

## Abbreviations

FFA2R: Free fatty acid receptor 2
DAMPs: Danger-associated molecular patterns
PAMPs: Pathogen-associated molecular patterns
GPCR: G protein-coupled receptor
FFAR: Free fatty acid receptor
SCFA: Short chain fatty acid
BHB: β-hydroxybutyrate
CTAB: Hexadecyltrimethylammonium bromide
BSA: Bovine serum albumin
HRP: Horseradish peroxidase
SOD: Superoxide dismutase
KRG: Krebs-Ringer Glucose phosphate buffer
DMEM: Dulbecco’s modified eagle medium
TR-FRET: Time-resolved fluorescence resonance energy transfer
CL: Chemiluminescence
ROS: Reactive oxygen species
NET: Neutrophil extracellular trap

## Acknowledgement

This work was supported by the Swedish Research Council, the Swedish government under the ALF-agreement, the Rune and Ulla Amlöv Foundation (JM), the Gothenburg society of Medicine (JM), the Clas Groschinskys Memorial Fund (MS M21146), the Åke Wiberg Foundation (MS M21-0025), the Sahlgrenska International Starting Grant (MS), the Magnus Bergwall Foundation (MS 2021-04110) and the Ingabritt and Arne Lundberg Foundation.

## Authors’ contributions

HF, CD and JM designed and oversaw all the aspects of the study. JM, MS, SL, MV, LZ and LB performed and analyzed the experiments with input from HF, XX and CD. HF and SG wrote the first draft. All authors revised the manuscript and approved the final version prior submission.

## Conflict-of-Interest Disclosure

The authors declare no conflict of interest. All authors fulfill the authorship criteria and declare no competing interests. Saray Gutierrez is an employee of AstraZeneca.

## Notes

### Competing Interest Statement

The authors have declared no competing interest.

